# Glucose increases the lifespan of post-reproductive *C. elegans* independently of FOXO

**DOI:** 10.1101/347435

**Authors:** Wang Lei, Caroline Beaudoin-Chabot, Guillaume Thibault

**Affiliations:** School of Biological Sciences, Nanyang Technological University, Singapore, 637551

**Author notes:** Correspondence to: Guillaume Thibault.

## Abstract

Aging is one of the most critical risk factors for the development of metabolic syndromes^1^. Prominent metabolic diseases, namely type 2 diabetes and insulin resistance, have a strong association with endoplasmic reticulum (ER) stress^2^. Upon ER stress, the unfolded protein response (UPR) is activated to limit cellular damage by adapting to stress conditions and restoring ER homeostasis^3,4^. However, adaptive genes upregulated from the UPR tend to decrease with age^5^. Although stress resistance correlates with increased longevity in a variety of model organisms, the links between the UPR, ER stress resistance, and longevity remain poorly understood. Here, we show that supplementing bacteria diet with 2% glucose (high glucose diet, HGD) in post-reproductive 7-day-old (7DO) *C. elegans* significantly extend their lifespan in contrast to shortening the lifespan of reproductive 3-day-old (3DO) animals. The insulin-IGF receptor DAF-2 and its immediate downstream target, phosphoinositide 3-kinase (PI3K) AGE-1, were found to be critical factors in extending the lifespan of 7DO worms on HGD. The downstream transcription factor forkhead box O (FOXO) DAF-16 did not extend the lifespan of 7DO worms on HGD in contrast of its previously reported role in modulating lifespan of 3DO worms^6^. Furthermore, we identified that UPR activation through the highly conserved ATF-6 and PEK-1 sensors significantly extended the longevity of 7DO worms on HGD but not through the IRE-1 sensor. Our results demonstrate that HGD extends lifespan of post-reproductive worms in a UPR-dependent manner but independently of FOXO. Based on these observations, we hypothesise that HGD activates the otherwise quiescent UPR in aged worms to overcome age-related stress and to restore ER homeostasis. In contrast, young adult animals subjected to HGD leads to unresolved ER stress, conversely leading to a deleterious stress response.

## INTRODUCTION

Diets of industrialised countries are enriched in processed carbohydrates or sugars which are rapidly converted into glucose, thus raising blood glucose levels very quickly^7^. High blood glucose level is strongly associated with metabolic syndrome such as type II diabetes (T2D) and cardiovascular diseases consequently affecting lifespan^8,9^. First shown in young *C. elegans*^6,10^ and subsequently in other model organisms, high-glucose diet (HGD) is sufficient to shorten lifespan^11,12^. In contrast, calorie restriction without malnutrition was initially shown in rodents and later in other model organisms including *C. elegans* to increase lifespan^10,13^. From these observations, the highly conserved insulin/IGF-1-signalling and TOR signalling pathways have been identified to modulate lifespan in several organisms^14–17^. However, the effect of HGD on the lifespan of post-reproductive animals remains to be addressed as prior observations on diet-related modulation of lifespan has been limited to young adult animals.

Aging is one of the most critical risk factors for the development of metabolic syndromes. Both T2D and insulin resistance have a strong association with endoplasmic reticulum (ER) stress. Upon ER stress, the unfolded protein response (UPR) is activated to limit cellular damage by adapting to stress conditions and restoring ER homeostasis. It activates multiple intracellular signalling pathways designed to restore ER and subsequently cellular homeostasis. The UPR programme results in a transient decrease of translation, enhanced degradation of misfolded proteins, and increased levels of ER-resident molecular chaperones^18^. However, the activation from genes through the UPR tend to decrease while the incidence of developing metabolic syndromes increases with age^5^. Although stress resistance correlates with increased longevity in a variety of model organisms, the links between the UPR, ER stress resistance and longevity remain poorly understood.

Evidence in pancreatic β cells showed that regulation of insulin secretion is dependent on PERK, one of the three sensors of ER stress. Coincidentally, specific mutations in PERK cause Wolcott-Rallison syndrome, which is characterised by early-onset diabetes mellitus^19^. Similarly, alternative ER stress sensing IRE1/XBP1 axis is required for glucose-induced insulin synthesis in pancreatic β cells^20,21^. Both PERK and IRE1 sensors are activated from ER stress as part of the UPR to restore ER homeostasis. Despite the strong association between ER stress and diabetes, the mechanism by which PERK regulates diabetes-related genes and the role of ER stress in the development of the disease are still largely unknown. Nevertheless, the failure of the UPR to restore homeostasis can become harmful to pancreatic β cells by eventually initiating apoptosis, thereby reducing insulin secretion, and contributing to T2D pathogenesis^22^.

Aging is also characterised by a gradual degeneration of physiological capacities and the diminished ability to cope with environmental stresses, which in turn leads to a heightened susceptibility to age-related diseases^23^. Several studies correlate aging to oxidative damage incurred by reactive oxygen species (ROS)^24^. These reactive compounds cause damage within the cell such as changing the properties of lipid membranes, cross-linking cellular proteins and causing mutagenic changes to DNA. Oxidative and ER stress intersect as some forms of ROS can activate the UPR^25,26^. To counteract aging, The UPR mitigates cellular damage brought about by stress conditions^27^ although the programme tends to decrease with age^5,28^. A similar decline has been observed for other homeostatic cellular stress responses^29,30^.

In *C. elegans*, when *daf-2* (insulin/IGF receptor) and *daf-16* (FOXO) mutants are challenged with HGD, it results in extended and shortened lifespans, respectively^6^. However, these assays were carried out in young adults. To assess the effect of HGD on post-reproductive organisms, we exposed 7-day-old (7DO) to HGD. To our surprise, 7DO worms subjected to HGD had a significantly longer lifespan compared to those fed with normal diet while 1- and 3-day-old (3DO) worms on HGD had significantly shorter lifespans. Unexpectedly, the lifespan extension observed in 7DO worms exposed to HGD was not dependent but was instead modulated by the PEK-1 and ATF-6 UPR sensors in post-reproductive worms.

## RESULTS

### High glucose diet extends lifespan of post-reproductive adult worms

The shortening of lifespan from high glucose diet in *C. elegans* as well as other model organisms is well documented^6,10–12^. However, little is known on the effect of dietary glucose on post-reproductive animals. To compare the effect of high glucose diet (HGD) on the lifespan of young adults and post-reproductive worms, wild type (WT) *C. elegans* of 1-, 3-, and 7-day-old were fed a diet of UV-killed *E. coli* OP50 bacteria supplemented with 2% glucose. As previously reported^6^, HGD significantly shortened the lifespan of 1- and 3-day-old worms compared to normal diet (ND) (Fig. 1a, Extended Data Table 1) and compromised the development of 1-day-old worms^31^ (Extended Data Fig. 1a). In contrast, 7-day-old (7DO) worms on HGD significantly extended their lifespan compared to ND (Fig. 1b, Extended Data Table 1). As pharyngeal pumping rate reduces with age^32^ and with HGD^31^, we asked if glucose reduces bacteria intake which could result in calorie restriction-induced longevity^10^. Pumping rate of 7DO worms subjected to 24h HGD was similar to normal diet (ND) (Fig. 1c). Subjecting 7DO worms to 4h starvation significantly increased pumping rate as previously reported^33^. This suggests that food intake is not influenced by HGD in aged worms. To further assess if food intake is influenced by HGD in 7DO worms, we measured the change of OP50 density in liquid culture after 24h incubation (Fig. 1d). In contrast to pumping rate, HGD significantly decreased bacteria intake in 7DO WT worms. Similarly, the accumulation of *E. coli* expressing free GFP^34^ in the intestines of WT worms was decreased by 42% with 24h HGD compared to ND (Fig. 1e). From these findings, we reasoned that aged worms might require fewer bacteria due to the excess of calorie intake from HGD. To investigate this hypothesis, we first measured glucose levels in aged WT worms fed ND or HGD (Fig. 1f). As expected, HGD significantly increased the glucose level of 7DO worms compared to ND as previously reported for young worms on HGD^6^. As the accumulation of glucose in aged worms could subsequently be utilized for ATP production^35^, we measured the ATP levels of whole worms. Aged WT worms fed 24h HGD revealed a significantly elevated ATP level when compared to aged WT worms on ND (Fig. 1g). In addition, HGD leads to an increase in fatty acids as well as oogenesis, suggesting a surge in metabolic activities (Extended Data Fig. 1b-d). To assess the long term outcome of HGD, we measured the motility of 14-day-old WT worms fed 7 days on ND or HGD as calorie restriction reduces motion^36^. WT worms fed HGD were significantly more active when compared to ND while having the opposite effect on 3DO worms (Fig. 1h). Taken together, our findings suggest that 7DO worms live longer from the excess of energy derived from HGD.

**Fig. 1.**
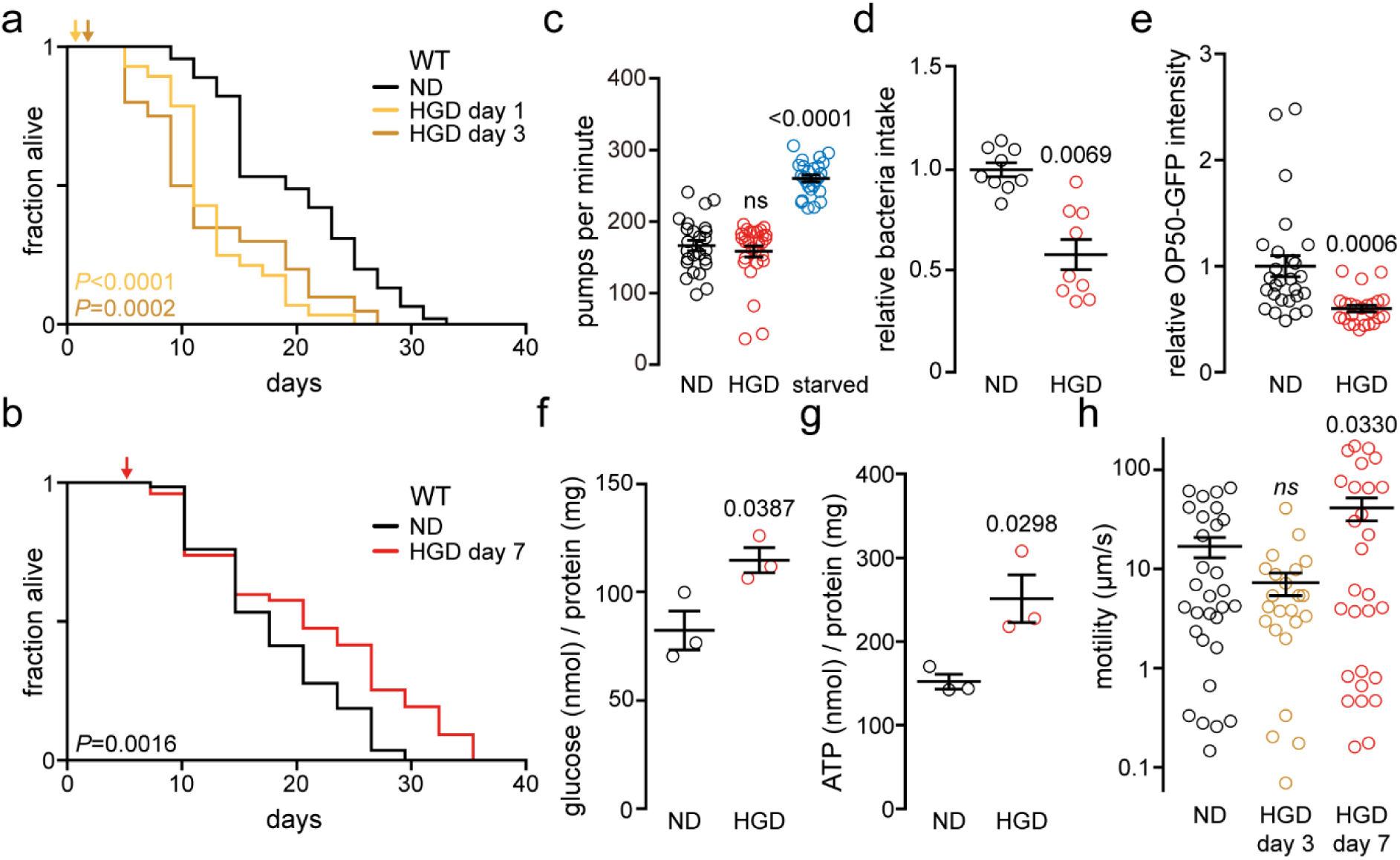
Glucose extends lifespan of post-reproductive but not of young adult worms. **a,** Lifespan assays of WT nematodes fed normal dead *E. coli* OP50 diet (ND) or ND supplemented with 2% glucose (high glucose diet, HGD) at 1- or 3-day old. **b,** Lifespan assay of 7-day-old WT worms treated as in **a**. **c,** Pharyngeal pumping rate in 7-day-old WT on 24h ND, HGD, or 4h starvation (ND, n=25; HGD, n=28; starved, n=25). **d,** Bacteria intake expressed relative to bacterial clearance within the WT ND interval of 7-day-old worms treated as in **c** (n=45). **e,** Quantification of confocal fluorescent microscopy images of GFP-expressing bacteria colonising WT worm intestine treated as in **c** (ND, n=27; HGD, n=24). **f,** Glucose levels inside of worms treated as in **c**. **g,** Total ATP concentration of worms treated as in **c**. **h,** Motility on bacteria-free nematode growth media (NGM) agar of worms treated as in **c** (ND, n=30; HGD day 3, n=23; HGD day 7, n=28). *P* values to ND of WT. *ns*, non-significant with *P*>0.05.

### Glucose-induced longevity of aged worms works independently of DAF-16/FOXO

The highly conserved insulin/IGF-1 signalling (IIS) pathway modulates growth, differentiation, and metabolism from nutrient availability and fluctuations in environmental conditions^14–17,37^. As the activation of IIS pathway shorten lifespan of young adult worms fed HGD^6^, we asked if the pathway modulates the longevity of 7DO worms on HGD. We carried out the lifespan assay of long-lived 7DO *daf-2(loss-of-function*; *lof)*^14^ and *age-1(lof)* animals subjected to HGD (Fig. 2a, b, Extended Data Table 1). DAF-2 and AGE-1 are the orthologues of the insulin-like growth factor 1 (IGF-1) receptor and the phosphoinositide 3-kinase (PI3K), respectively. AGE-1 is activated downstream of DAF-2 receptor^38^. Similar to WT worms, HGD significantly extended the lifespan of 7DO *daf-2(lof)* worms compared to ND. However, the difference was not as pronounced as in WT worms (Fig. 1a, Extended Data Table 1). In contrast, there was no significant difference in the lifespan of 7DO *age-1(lof)* on HGD compared to ND. To better understand this discrepancy in the effect of HGD on the lifespan of *daf-2(lof)* and *age-1(lof)*, we sought to avoid the manual selection of normally developing *daf-2(lof)* worms (~75%) from dauer stage worms (~25%) induced at 20°C^39^. Thus, we incubated larva stage 1 (L1) *daf-2(lof)* worms at 16°C which yielded ~98% of normally developing *daf-2(lof)* worms. We subsequently conducted the lifespan assay at 20°C. HGD did not extend the lifespan of *daf-2(lof)* worms compared to control using this unconventional temperature shift (Extended Data Fig. 2, Extended Data Table 1). It should be noted that WT worms were treated as *daf-2(lof)* worms and exhibited the consistent lifespan extension upon HGD as shown for WT worms constantly incubated at 20°C (Fig. 1a, Extended Data Fig. 2 and Table 1). Together, these findings suggest that DAF-2 and AGE-1 play a role in extending the lifespan of 7-day-old worms fed with HGD. This is in contrast to the role of DAF-2 and AGE-1 in reducing the lifespan of young adult worms fed HGD^6^.

**Fig. 2.**
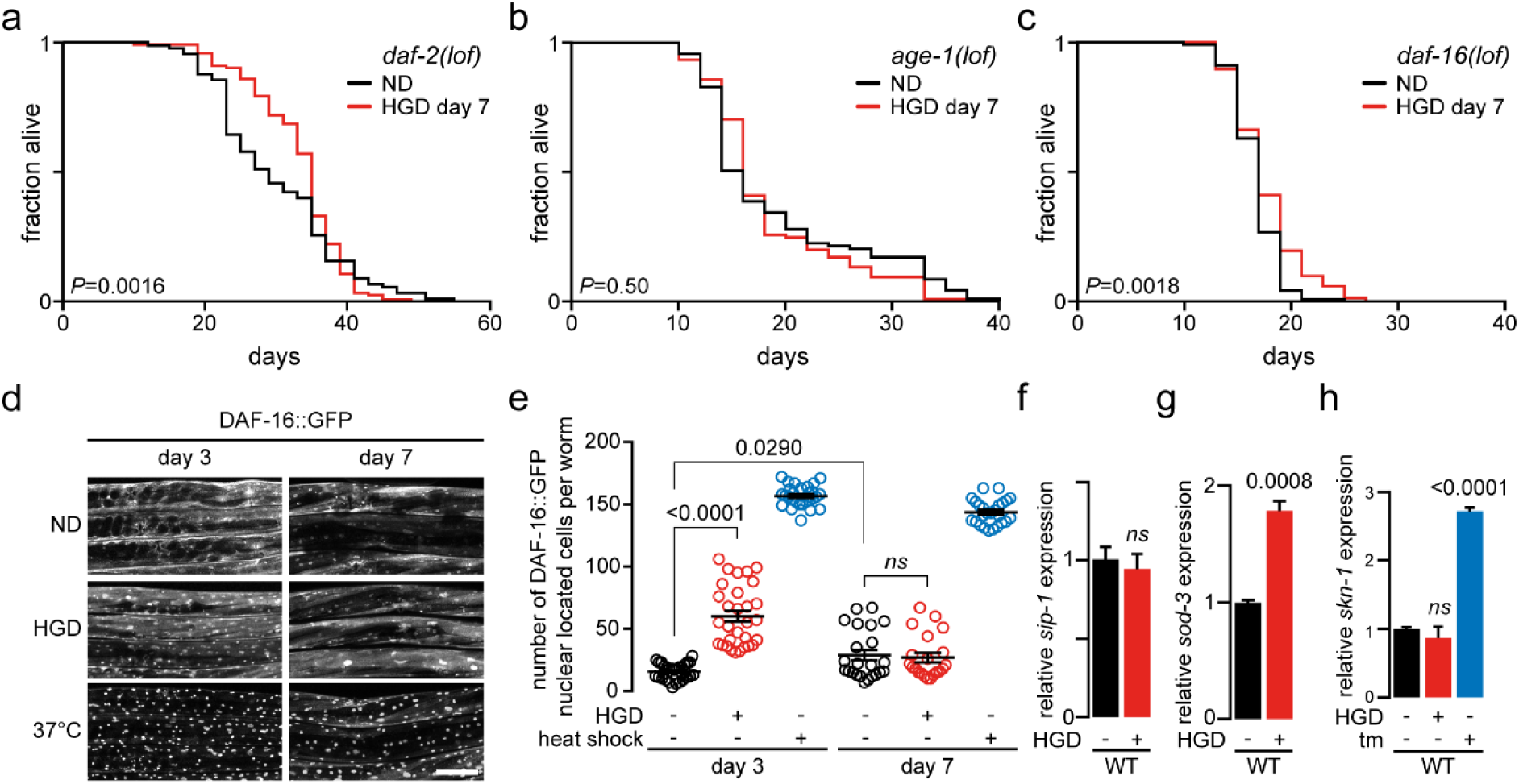
Glucose extends lifespan in aged worms independently of the DAF-16 (FOXO). **a-c,** Lifespan assays of *daf-2(lof)* (**a**), *age-1(lof)* (**b**), and *daf-16(lof)* (**c**) animals fed normal diets (ND) or high glucose diet (HGD) from 7-day-old. **d,** Representative images of DAF-16::GFP localization in 3- and 7-day-old worms on 16h ND, HGD or 1h heat shock at 37°C. **e,** Quantification of **d** (n=31, n=30, n=28 for ND, HGD, heat shock, respectively, for day 3; n=23, n=21, n=25 for ND, HGD, heat shock, respectively, for day 7). **f, g,** qPCR comparing expression of genes *sip-1* (**f**) and *sod-3* (**g**) in 7-day-old WT on 24h ND or HGD. **h,** qPCR comparing expression of genes *skn-1* in 7-day-old WT worms on 24h ND, HGD or 4h tunicamycin (Tm). *P* values to ND of WT. *ns*, non-significant with *P*>0.05.

To further investigate the role of the IIS pathway in modulating the lifespan of post-reproductive worms on HGD, we carried out the lifespan assay on *daf-16(lof)* animals. DAF-16 is the orthologue of transcription factor forkhead box O (FOXO) which is inhibited from DAF-2 and AGE-1 activation^38^. Seven-day-old *daf-16(lof)* lived significantly longer on HGD compared to normal bacterial diet, and this effect was not significantly different from WT on HGD (Fig. 2c, Extended Data Table 1). This finding suggests that DAF-16 is not involved in extending the lifespan of aged worms on HGD. In contrast, glucose has been reported to downregulate DAF-16/FOXO activity in young adult worms (3-day-old, 3DO) resulting in shorter lifespan^6^.

To ensure DAF-16 is inducible in 7DO worms as in young adults, we visualised DAF-16::GFP localisation in 3- and 7-day old worms (Fig. 2d, e). In 3-day-old worms fed 16h HGD, we observed a significant increase in nuclear DAF-16::GFP localisation, indicating a HGD-induced translation of DAF-16. In contrast, no significant activation of DAF-16::GFP was observed in 7DO worms on HGD. As expected, DAF-16::GFP strongly localised to the nucleus in 3- and 7-day-old worms upon heat shock^40^, indicating DAF-16 activation is not diminished with age. Next, we assessed the expression levels of DAF-16 target genes *sip-1* and *sod-3* by quantitative PCR (qPCR). There was no significant change in the expression of *sip-1* upon HGD (Fig. 2f), suggesting the activity of DAF-16 is unchanged. In contrast, expression of *sip-1* was significantly increased in 3DO worms fed HGD (Extended Data Fig. 4c). The expression of *sod-3* was higher in WT worms on HGD, reflecting an increase of glucose-induced oxidative stress^41^ (Fig. 2g). We also monitored the expression levels of the transcription factor *skn-1* which could potentially extend *C. elegans* lifespan and is inhibited through the DAF-2/AGE-1 axes in a similar fashion as DAF-16^38^. The drug tunicamycin (tm) was used to induce ER stress and consequently upregulates *skn-1* through the activation of the unfolded protein response (UPR)^42^. As DAF-2 is activated from HGD, no significant change was observed in *skn-1* levels in 7DO worms on HGD compared to ND (Fig. 2h). Together, the data indicate that the IIS pathway remains intact and functional in aged worms. Accordingly, the HGD-induced longevity observed in aged worms is modulated independently from DAF-16 (FOXO).

### The unfolded protein response is required to extend lifespan of post-reproductive worms on glucose diet

As DAF-16 can be excluded, we turned our attention to other pathways that influence longevity. As HGD induces ER stress, we monitored the expression of genes *cht-1*, *hsp-4*, and *F40F12.7* upregulated from the UPR branches ATF-6, IRE-1, and PEK-1, respectively^43^ (Extended Data Fig. 3). The UPR is activated to limit cellular damage by adapting to stress conditions and to re-establish ER homeostasis while an acute UPR activation drives pro-apoptotic pathways^3^. Feeding HGD to 3- and 7-day-old WT worms induced ER stress with a significant activation of the UPR programme through all three sensors compared to ND (Fig. 3a). However, HGD-induced upregulation of hsp-4 through the IRE-1 branch was significant lower in 7DO compared to 3DO worms which is consistent with previous findings^5^. On the other hand, there was a significant upregulation of *cht-1* and *F40F12.7* from HGD in 7DO compared to 3DO worms. This suggests that ATF-6 and PEK-1 might compensate for the weak activation of IRE-1 in 7DO animals, consequently playing an important role in maintaining ER homeostasis in aging animals. To explore the role of the UPR in modulating longevity of post-reproductive worms subjected to HGD, we monitored the lifespan of *atf-6(lof)*, *ire-1(lof)*, and *pek-1(lof)* animals. HGD significantly extended the lifespan of 7-day-old *ire-1(lof)* but not *atf-6(lof)* and *pek-1(lof)* when compared to ND, indicating that both ATF-6 and PEK-1 play a role in extending the longevity of aged worms on HGD (Fig. 3b-d, Extended Data Table 1). Similar to WT animals, we carried out the same experiments to ensure that lifespan extension was not dependent on DAF-16 and SKN-1 (Extended Data Fig. 4) as well as a consequence of calorie restriction for the UPR mutants (Extended Data Fig. 5). Interestingly, PEK-1 was found to be critical for organismal development while ATF-6 is dispensable in *C. elegans* on HGD from larva stage 1 (Extended Data Fig. 5e).

**Fig. 3.**
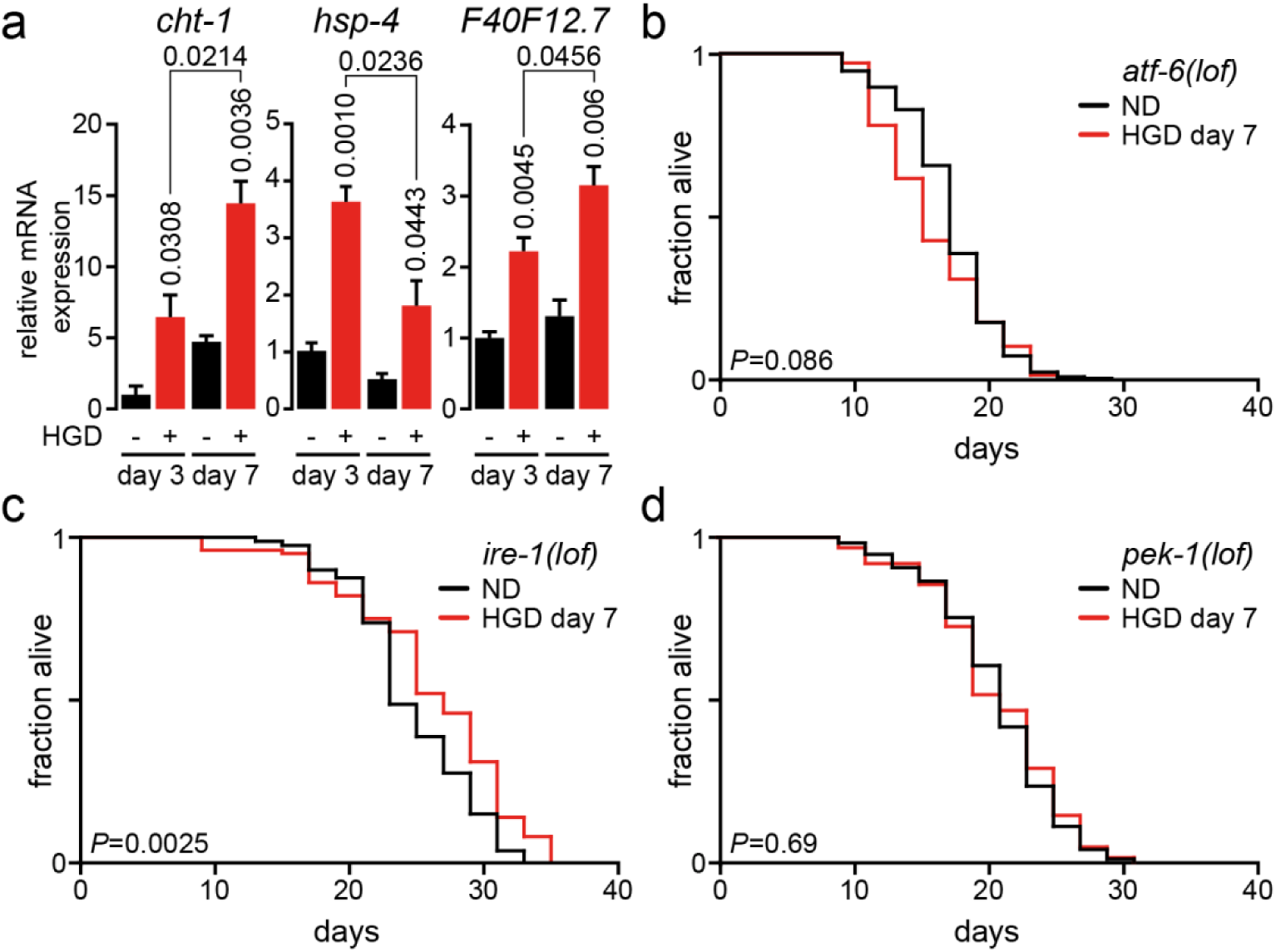
ATF-6 and PEK-1 are essential to extend lifespan of aged worms exposed to HGD. **a,** qPCR comparing expression of genes *cht-1*, *hsp-4*, and *F40F12.7* induced from UPR branches ATF-6, IRE-1, and PEK-1, respectively, in 3- and 7-day-old WT worms fed 24h normal diet (ND) or high glucose diet (HGD). **b-d,** Lifespan assay of 7-day-old *atf-6(lof)* (**b**), *ire-1(lof)* (**c**), and *pek-1(lof)* (**d**) worm mutants fed ND or HGD. *P* values to respective ND of WT.

### HGD-activated UPR regulates longevity and metabolism of worms by affecting multiple tissues

To further dissect how the UPR modulates HGD-induced longevity of post-reproductive worms, we carried out an RNAi screen using the glucose inducible reporter strain *far-3p::GFP*^44^. We screened RNAi clones of common genes to the reported suppressors and enhancers of glucose-induced *far-3p::GFP* and UPR-upregulated genes^43,45^ (Fig. 4a). Major UPR related genes *atf-5*, *atf-6*, *ire-1*, and *pek-1* were included to the screen. Seven-day-old worms fed on normal diet were transferred to a 24 h HGD in combination with RNAi clones. Out of the 216 clones screened, GFP fluorescence was normalised to the RNAi empty vector (pL4440) resulting in one candidate enhancing and 173 suppressing the GFP reporter at more than 1.5 fold difference (Extended Data Table 2). Notable, *atf-6* and *ire-1* RNAi significantly decreased the GFP signal, confirming the important role of the UPR in glucose-derived metabolism. Next, we performed functional annotation of the significant UPR-related suppressors of glucose-induced FAR-3 using the gene ontology (GO) and tissue enrichment tool DAVID (Fig. 4b-f and Extended Data Table 3). Ninety seven of 168 genes were enriched in the intestine as this organ digest and metabolise glucose^46^ in addition to playing an important role in increasing lifespan through the UPR^5^. Interestingly, 11 genes playing a role in adult lifespan were enriched in the intestine, suggesting some of these genes might be key regulators of HGD-induced longevity of post-reproductive worms. Immune response and lipid metabolic regulatory genes were enriched in the intestine, outer labial sensillum, and PVD neurons, suggesting an important role of UPR-modulated genes. Both insulin-like genes *ins-2* and *ins-5* were enriched in the head muscle, pointing to the important role of IIS pathway in glucose metabolism. Together, the screening suggest that the UPR plays an important role to maintain ER homeostasis through high metabolism in the intestine.

**Fig. 4.**
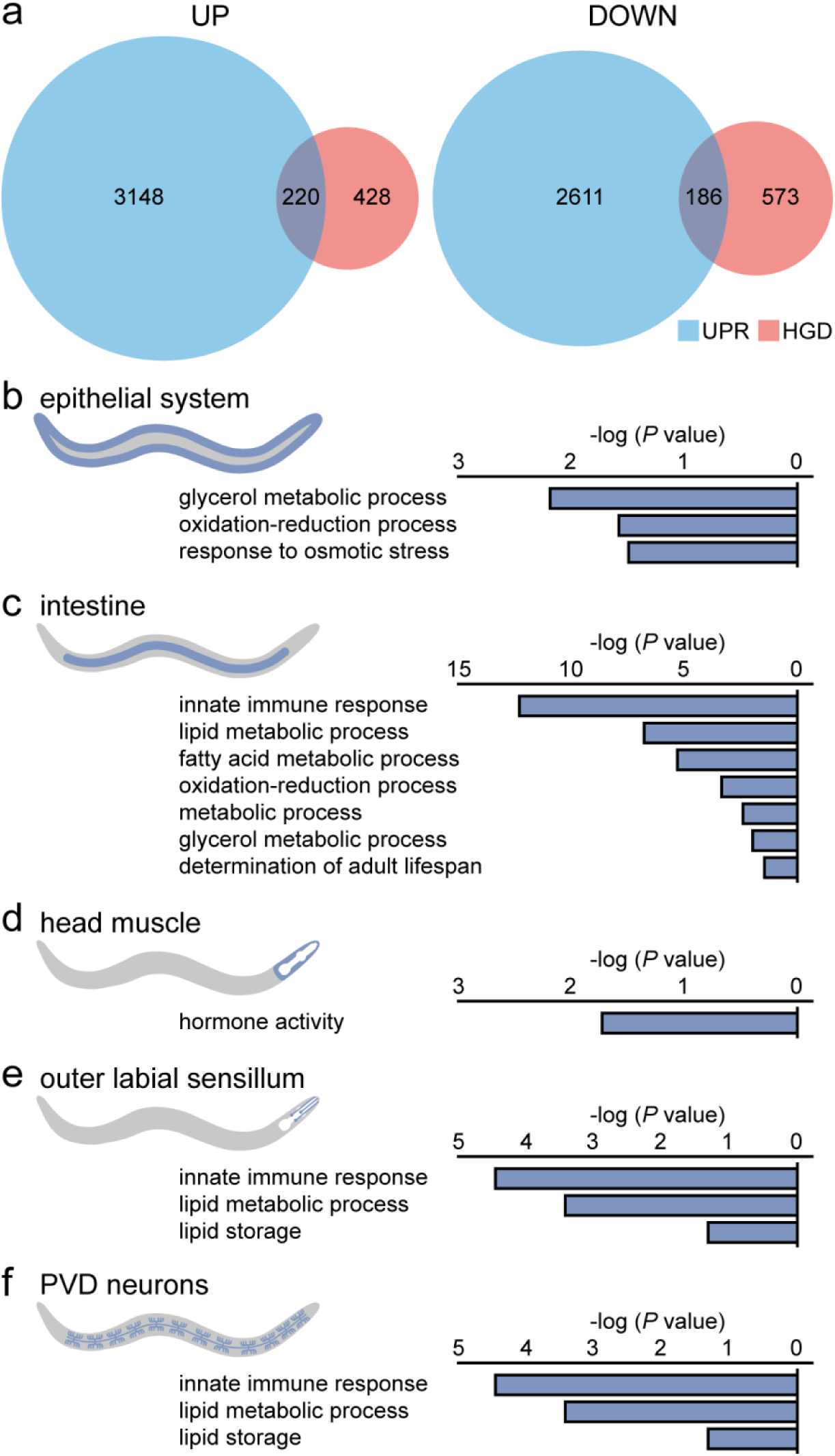
A subset of genes is modulated from HGD and the UPR. **a,** Venn diagram representation of upregulated and downregulated genes at a minimum of 1.5 fold by the unfolded protein response (UPR) and genes ND worms and HGD fed worms**. b-f,** tissue enrichment and gene ontology analysis for target genes filtered by glucose *in vivo* reporter strain at a minimum of 1.5 fold in fluorescence intensity.

## DISCUSSION

Protein quality control is essential to maintain cellular homeostasis where surveillance in the ER is provided by the UPR^4^. Defects in the pathway compromise the proteostasis network in the ER and promote the accumulation of damaged and misfolded proteins that might result in aggregate accumulation which is further accelerated during aging^47^. ER-localised protein disulfide isomerases (PDIs), chaperone calreticulin (CRT-1), and HSP-4 paralogue, HSP-3 (Hsp70 orthologue), were reported to decline by about two-fold throughout the course of the *C. elegans* lifespan. Similarly, ER stress-induced upregulation of HSP-4 through the UPR branch IRE-1 is dramatically attenuated during aging^5^. On the other hand, the downstream target mRNA of the UPR sensor IRE-1, *xbp-1*, has been shown to act in concert with DAF-16 to enhance ER stress resistance and thus longevity of IIS mutant worms^48^. Consequently, the intervention of the UPR in maintaining ER and cellular homeostasis is critical to extend lifespan. However, the fitness of the ATF-6 and PEK-1 branches of the UPR is largely unknown during aging. Here, our data suggest that the decline of IRE-1-mediated UPR responsiveness, in post-reproductive worms, compromises cellular homeostasis^47^, and ER stress agents such as high glucose is needed to sufficiently activate the highly conserved UPR sensors ATF-6 and PEK-1 to promote longevity (Fig. 5). Future studies should identify drugs that will specifically activate the ATF-6 and PEK-1 pathways without leading to an acute UPR activation. Ideally, the fitness of the UPR should be kept constant through life and especially during old age to prevent age-related accumulation of cellular damage^49^ and consequently premature death.

**Fig. 5.**
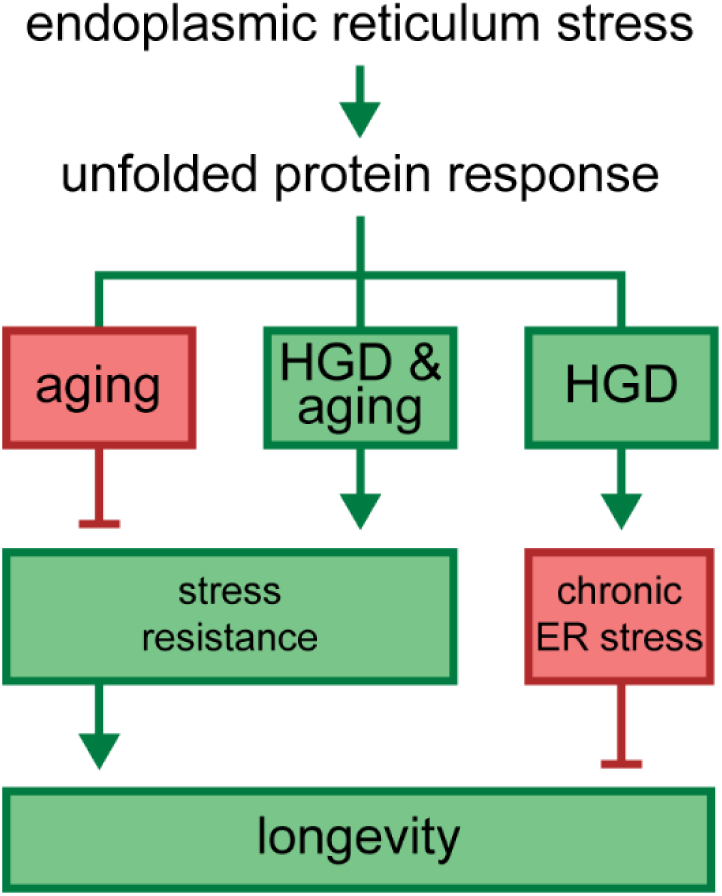
HGD promotes longevity in post-reproductive worms. Proposed model of age-dependent high glucose diet (HGD)-induced endoplasmic reticulum stress.

Aging is defined as a progressive loss of physiological integrity including reduced fertility, decreased protein homeostasis, cellular senescence, genomic instability and disrupted metabolic homeostasis^50^. Particularly relevant, sterility has been shown to promote longevity in a DAF-16-dependent manner using the germ-cell loss *C. elegans* mutant *glp-1(lof)*^51^. Endocrine signalling and multiple transcriptional pathways were subsequently reported to increase lifespan in gonad-ablated animals^52^. Additionally, an increase in insulin signalling, through HGD, promotes germline proliferation^53^. Thus, germline proliferation in young adult worms combined with HGD could possibly yield a synergetic effect, resulting in accelerated aging. On the other hand, HGD starting at the post-reproductive life stage might stimulate age-attenuated cellular stress responses, consequently extending lifespan through the restoration of cellular homeostasis. The severe decline in proteostasis correlates with the termination of germline proliferation^54^. Our study points to a specific life stage from which it could be beneficial to artificially promote proteostasis and consequently counteract the effects of aging. The timing for intervention might be critical, as it could exacerbate aging if done too early. It remains to be seen if therapeutic treatment to activate the UPR in post-reproductive mammals will correlate with an extension of lifespan through the clearance of intracellular damages.

Extended lifespan is coupled with an increase in motility and energy metabolism^55,56^, indicative of healthy aging. We found that post-reproductive worms were metabolically more active on HGD as they displayed higher motility, an increase in total ATP and oogenesis, thereby suggesting healthier aging compared to worms fed with normal diet. As the proteostasis network decline during aging, proper digestion of bacteria including pharyngeal grinding, intestinal lysozymes, intestinal sapsins and amebapores might be impaired in aging *C. elegans*^46^. Evidence also suggests that the transcriptome and proteome relating to glucose metabolic pathways differ from young to post-reproductive worms^47,57^. Similarly in aging mice, metabolites derived from glucose, fatty acids, and amino acids decline dramatically upon aging^58^. It could be hypothesised that higher levels of HGD-induced insulin production might be required to stimulate the IIS pathway while its responsiveness might decline upon aging. Consequently, it is reasonable to conceive that supplementing the diet with glucose will restore healthy calorie intake in aged animals while becoming excessive combined with well digested bacteria in young animals. However, it remains to be determined weather or not calorie restriction initiated at the post-reproductive stage will result in lifespan extension as demonstrated in young animals^10^.

The highly conserved insulin signalling pathway promotes lifespan extension when attenuated or muted in *C. elegans* and other multicellular organisms^59^. Ablation of *age-1* or *daf-2* genes double the lifespan of *C. elegans* while the absence of the FOXO transcription factor DAF-16 dramatically accelerates tissue aging and reduces longevity^14,40,60^. HGD is sufficient to reduce longevity by downregulating DAF-16 activity through DAF-2 activation in young adults^6^. Similarly, calorie restriction increases lifespan by upregulating DAF-16 activity, increasing cellular stress surveillance^61,62^. Surprisingly, our findings suggest that while the fitness of DAF-16 is not attenuated during aging, it does not play a role in promoting the lifespan of post-reproductive worms subjected to HGD. It is reasonable to conclude that DAF-16 is inactivated through DAF-2 as both DAF-2 and AGE-1 are required to extend lifespan in post-reproductive worms. Perhaps, they both function in promoting glucose uptake and metabolism in order to extend lifespan of aged animals^63^. In addition, DAF-2 and AGE-1 could possibly be required through crosstalk with the UPR programme as the IRE-1/XBP-1 axis has been previously reported to compensate for the ablation of the IIS pathway^48^. The interdependence of the IIS pathway with the UPR branches ATF-6 and PEK-1 needs to be explored in the future to fully grasp their role in HGD-induced longevity of post-reproductive worms.

## ACKNOWLEDGEMENTS

We are grateful to our colleagues Drs Valerie Lin Chun Ling and I-Hsin Su as well as Thibault’s lab members Xiu Hui Fun, Nurulain Ho, and Peter Jr. Shyu for critical reading of the manuscript. We thank Dongyeop Lee and Dr Seung-Jae V. Lee for generously sharing the glucose-feeding RNA sequencing data and glucose reporter strain *far-3p::GFP(bc14852)*^44^. Some strains were provided by the *Caenorhabditis* Genetics Center, which is funded by NIH Office of Research Infrastructure Programs (P40 OD010440). This work was supported by the Nanyang Assistant Professorship programme from Nanyang Technological University and the Singapore Ministry of Education Academic Research Fund Tier 1 (2016-T1-001-066).

## AUTHOR CONTRIBUTIONS

W.L. and G.T. designed the studies, W.L. and C.B.C. performed all the experiments. W.L. and G.T. wrote the manuscript with input from all the authors.

## COMPETING INTERESTS

The authors declare that they have no conflict of interest.

## METHODS

### Statistics

Error bars indicate standard error of the mean (SEM), calculated from at least three biological replicates, unless otherwise indicated. *P* values were calculated using one-way ANOVA with Tukey’s test or log-rank test for lifespan, unless otherwise indicated and reported as *P* values with 4 significant digits in the figures. All statistical tests were performed using GraphPad Prism 7 software (GraphPad Software, Inc., San Diego, CA).

### *C. elegans* strains, RNAi constructs, and bacterial strains

All strains were grown at 20°C using standard *C. elegans* methods as previously described^64,65^. Nematode growth media (NGM) agar plates were seeded with *Escherichia coli* strain OP50 (UV-killed bacteria for lifespan assays, ultrasound-killed bacteria for liquid culture) for normal growth and with HT115 bacteria for RNAi feeding. NGM agar plates were supplemented with 2% glucose (high glucose diet, HGD) when indicated. RNAi feeding was performed as previously described^66^, and the RNAi library was obtained from the Fire lab^67^. The plasmids were verified by sequencing. *C. elegans* strains wild type N2, *atf-5(ok576), atf-6(ok551), ire-1(ok799), pek-1(ok275), pmt-2(vc1952), daf-2(e1370), daf-16(mu86)*, *hsp4::GFP*(*sj4005*), *daf-16::GFP(tj356), far-3p::GFP(bc14852)* and bacteria strains *OP50* as well as *OP50*-*GFP* were gifted from the *Caenorhaditis* Genetics Center.

### Lifespan assay

Lifespan assays were performed at 20°C as previously described^68^. Animals were transferred to NGM plates containing 2% glucose at 1-, 3-, or 7-day-old from hatching when indicated. Day 1 was defined as the egg hatching event. Pyrimidine analogue 5-fluoro-2’-deoxyuridine (FUdR, Sigma; St. Louis) was added at 50 µM to L3/L4 stage worms to prevent development of progeny. Adults were scored manually as dead or alive every 2-3 days. Nematodes that ceased pharyngeal pumping and had no response to gentle stimulation were recorded as dead. Worms that were male or dead due to desiccation were excluded from the analysis.

### Quantitative real-time PCR

Quantitative real-time PCR (qPCR) screen was carried out as previously described with modifications^45^. Approximately 1,000 synchronised worms were grown on NGM plates and harvested after 3 or 7 days. Worms were lysed with a motorised pestle homogeniser. Total RNA was isolated using TRIzol reagent (Thermo Fisher, Waltham, MA). Contaminant DNA was removed from samples with TURBO DNase (Thermo Fisher) following the manufacturer’s protocol. Complementary DNA (cDNA) was synthesised from total RNA using RevertAid Reverse Transcriptase (Thermo Fisher, Waltham, MA) following the manufacturer’s protocol. qPCR was performed with SYBR Green (Qiagen, CA, USA) following manufacturer’s protocol using a CFX-96 Real-time PCR system (Bio-Rad, Hercules, CA, USA). Thirty nanograms of cDNA and 50 nM of the paired primer mix for target genes were used for each reaction. Relative mRNA was normalised to the housekeeping gene *act-1*. Primers for qPCR are available upon request.

### Fatty acids analysis

Approximately 10,000 day 7 worms were harvested and washed thoroughly with M9 buffer and lyophilised overnight (VirTis, Warminster, PA). Fatty acids were esterified to fatty acid methyl esters (FAME) with 300 µl of 1.25 M HCl-Methanol for 1h at 80°C. FAMEs were extracted three times with 1 ml of hexane. Combined extracts were dried under nitrogen stream and resuspended in 20 µl hexane. FAMEs were separated by gas chromatography with flame ionization detector (GC-FID) (GC-2014; Shimadzu, Kyoto, Japan) using an ULBON HR-SS-10 50 m × 0.25 mm column (Shinwa, Tokyo, Japan). Supelco 37 component FAME mix was used to identify corresponding fatty acids (Sigma-Aldrich, St. Louis, MO). Data was normalized using pentadecanoic acid (C15:0) internal standard and worm dry weight.

### Fluorescence microscopy

To quantify DAF-16::GFP localization and GFP expression, worms were immobilised with 25 mM tetramisole and mounted on 2% agarose pad. Images were captured using Zeiss LSM 800 confocal fluorescence microscope (Carl Zeiss AG, Oberkochen, Germany) with 20x HC PL objective lens wit excitation at 488 nm and emission at 550(13) nm. Two-micrometer Z-stack sections were merged and number of cells with nuclear-localized DAF-16::GFP was determined using Fiji imaging software.

### Pharyngeal pumping assay

Pharyngeal pumping was measured by monitoring the number of pharyngeal bulb contractions in a 60-second interval. Videos were recorded with a Dino-eye Eyepiece camera (Dino-eye, Taiwan) fitted onto a stereomicroscope (Nikon, Tokyo). Contractions were carefully monitored through videos at 0.4X playback speed. At least 20 worms were recorded for each group. Worms that exhibited under 5 contractions per 10 seconds were considered as not pumping and excluded from experiment.

### Quantification of bacteria intake

The consumption of *E. coli* OP50 by worms was assessed in 96-well plates as previously described^69^. Each well contained 4 to 10 7-day old worms and 3 mg/ml ultrasound-killed bacteria in 150 μl S-complete liquid medium supplemented with 50 µg/ml carbenicillin and 50 μM FUdR. After 24 hours incubation, plates were shaken at 200 rpm for 15 min and the absorbance at 600 nm was measured. The relative amount of bacteria consumed was obtained from the difference in the absorbance at 600 nm normalized to the number of worms. The same drug-supplemented media without worms was set as blank. At least 45 nematodes were recorded for each group. Bacteria intake was calculated from the difference in absorbance at over the 24h period and normalized to the number of worms.

The ingestion of *E. coli* OP50 GFP was measured as the fluorescence intensity *in vivo*. Briefly, worms were washed 3 times and resuspended in M9 buffer for 15 minutes to wash off any OP50-GFP that may have attached to the body wall. Worms were immobilised with 25 mM tetramisole and mounted on 2% agarose pad. Images were captured using Zeiss LSM 800 confocal microscope with 20x HC PL objective lens. Two micrometers Z-stacks sections were merged, and the total fluorescence intensity of each individual worm was quantified using Fiji imaging software. At least 20 nematodes were recorded for each group.

### Glucose level analysis

Glucose levels were measured as previously described^6^ with some modifications. Approximately 1,000 worms were harvested following 24h growth on NGM supplemented with 2% glucose when indicated. Pelleted worms were lysed by beat beating in RIPA buffer and spun down at 12,000 × *g* for 5 minutes, 4°C to remove the debris. The clarified lysate was transferred to a new tube and protein content was determined by bicinchoninic acid (BCA) assay. Trichloroacetic acid (TCA) was added to the remaining lysate to a final concentration of 10% and spun down at 12,000 × *g* for 5 minutes, 4°C. The supernatant was neutralised to pH 7 with 1 N NaOH containing 100 mM potassium phosphate and glucose concentration was measured using the Amplex Red Glucose/Glucose Oxidase kit (Thermo Fisher, Waltham, MA).

### ATP level analysis

ATP concentration was measured using the Promega ENLITEN ATP assay kit (Madison, WI) as previously described^70^. Worm samples were treated as the glucose level analysis up the step of neutralising the supernatant to pH 7. Protein concentration was determined using the Bicinchoninic Acid (BCA) Protein Assays kit following manufacturer’s protocol (Sigma; St. Louis). ATP concentration was determined following the instruction of ENLITEN ATP assay kit (Promega, Madison, WI).

### Measurement of body length

Body length was measured as previously described^71,72^ with some modifications. Worms were fed with normal or HGD upon hatching and harvested after 60 hours. They were then transferred to NGM plates without OP50. Images were taken using a Dino-eye Eyepiece camera fitted onto a stereomicroscope. The data was analysed using wrMTrck plugins in Fiji Image J.

### Computational analysis of glucose-induced UPR gene candidates

Microarray data of UPR-specific genes for lipid bilayer stress (*pmt-2* RNAi) and proteotoxic stress (tunicamycin) was adapted from our previous study ^45^. The list of genes regulated by 2% glucose diet supplementation was kindly provided by Dr. Seung-Jae V. Lee‘s lab^44^. Gene expression profiles that were either increased or decreased significantly by at least 1.5 fold were recorded. Venn diagram was generated based on Venny2.1.0^73^ and edited in Adobe Illustrator.

### RNAi screening

RNAi screen was carried out as previously described with modifications^74^. Briefly, synchronised L1 larval stage *far-3p::GFP* animals were grown NGM plates and harvested at 7-day-old. Five to ten M9-washed worms were seeded into 96-well microplates containing RNAi Bactria clones of previously identified HGD-modulated ^44^ and UPR-modulated genes^43,45^ in the absence of presence of 2% glucose. Fluorescence intensity was measured by Incell 2200 Analyzer (GE Healthcare Lifesciences) 24h post-seeding with excitation at 488 nm and emission at 515 nm. GFP fluorescence intensity was normalised to the number of worms per well. The genes with filtered by a minimum 0.5 fold change. The list of genes generated were further analysed by DAVID using the HGD-modulated ^44^ and UPR-modulated genes set as background^75,76^. The genes enriched from DAVID analysis were further analysed by Gene Set Enrichment Analysis in wormbase^77^.

## EXTENDED DATA FIGURES AND TABLES

**Extended Data Fig. 1.**
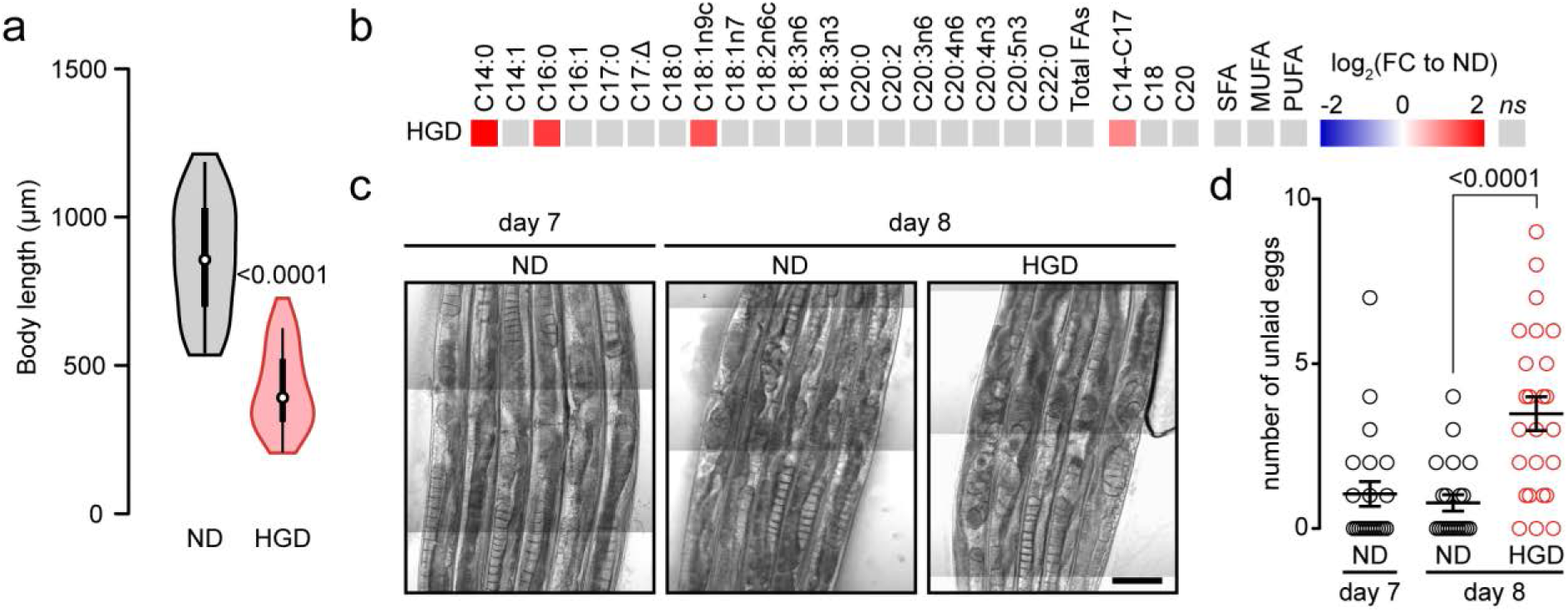
Seven-day-old WT worm is metabolically more active on high glucose diet. **a,** Body length distribution of WT worms fed normal diet (ND) or high glucose diet (HGD) at 48h from larva stage 1. **b,** Heat map of based 2 logarithmic fold changes (FC) in fatty acids (FAs), distribution of FA length, and saturation of 7-day-old worms on 24h HGD compared to WT ND. SFA, saturated fatty acid; MUFA, monounsaturated fatty acid; PUFA, polyunsaturated fatty acid. *ns*, non-significant with *P*>0.05. **c,** Representative images of unlaid eggs in 7DO WT worms on ND and after 24h ND or HGD. **d,** Number of unlaid eggs from **c** (n=22, n=22, n=25 for day 7 ND, day 8 ND, day 8 HGD, respectively).

**Extended Data Fig. 2.**
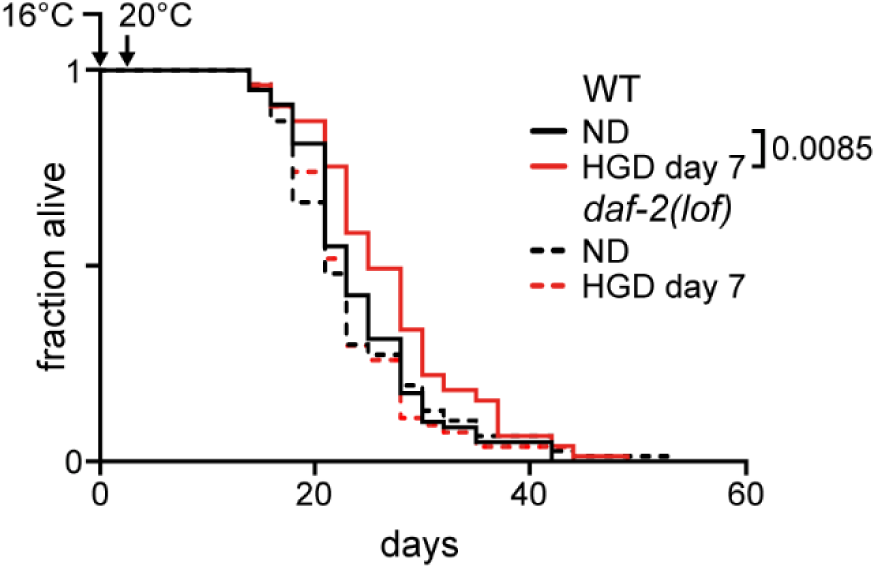
DAF-2 plays a role in extending lifespan of worms grown at 16°C during development. Lifespan assay of 7-day-old WT and *daf-2(lof)* animals fed normal diet (ND) or high glucose diet (HGD). Worms were grown at 16°C from larval stage 1 to 3 and dauer *daf-2(lof)* animals were discarded. Worms were subsequently grown at 20°C for the remaining of the assay. None reported *P* values were non-significant with *P*>0.05.

**Extended Data Fig. 3.**
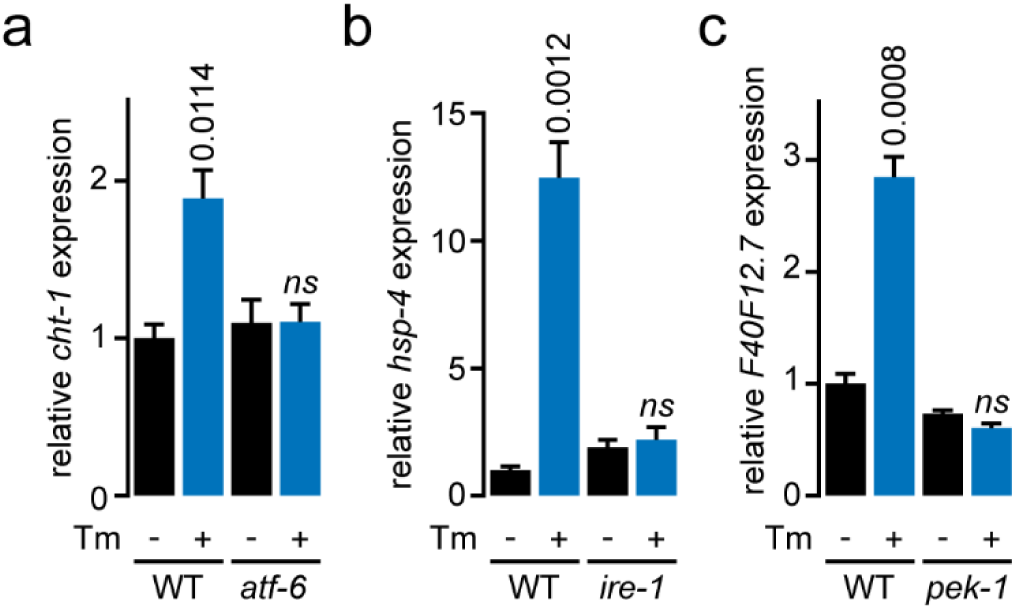
Validation of UPR target genes. **a**, qPCR comparing expression of genes *cht-1* in 3-day-old WT and *atf-6(lof)* worms fed normal diet (ND) or ND followed by 4h exposure to tunicamycin (Tm). **b**, qPCR comparing expression of genes *hsp-4* in 3-day-old WT and *ire-1(lof)* worms treated as in **a**. **c**, qPCR comparing expression of genes *F40F12.7* in 3-day-old WT and *pek-1(lof)* worms treated as in **a**. *P* values to respective WT or mutant worms without Tm. *ns*, non-significant with *P*>0.05.

**Extended Data Fig. 4.**
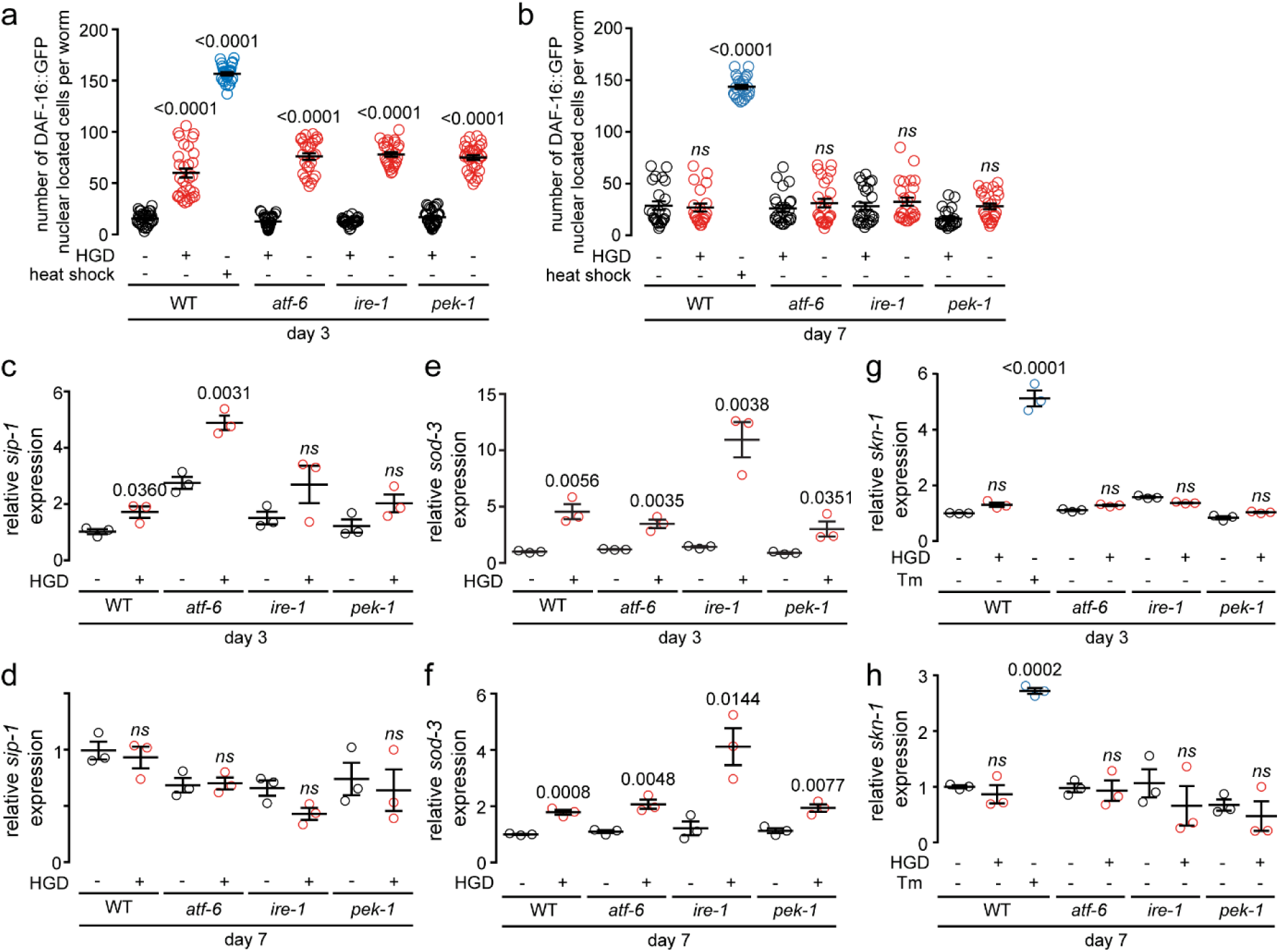
IIS pathway related genes expression. **a, b,** DAF-16::GFP localisation in 3-day-old (**a**) (n=31, n=30, n=28 for wild type worms on ND, HGD, heat shock condition, respectively; n=29, n=26 for *atf-6* mutant worms on ND, HGD respectively; n=30, n=27 for *ire-1* mutant on ND, HGD respectively; n=30, n=31 for *pek-1* mutant worm on ND, HGD respectively) and 7-day-old worms (**b**) on 24h normal diet (ND), high glucose diet (HGD) or subjected to 1h heat shock at 37°C (n=23, n=21, n=25for wild type worms on ND, HGD, heat shock condition, respectively; n=24, n=25 for *atf-6(lof)* on ND, HGD respectively; n=25, n=25 for *ire-1(lof)* on ND, HGD respectively; n=22, n=25 for *pek-1(lof)* on ND, HGD respectively). **c, d,** qPCR comparing expression of genes *sip-1* in 3-day-old (**c**) or 7-day-old (**d**) WT and UPR mutant worms fed 24h ND or HGD. **e, f,** qPCR comparing expression of gene *sod-3* in 3-day-old (**e**) or 7-day-old (**f**) WT and UPR mutant worms fed 24h ND or HGD. **g, h,** qPCR comparing expression of gene *skn-1* in 3-day-old (**g**) or 7-day-old (**h**) WT and UPR mutant worms fed 24h ND, HGD, or 4h tunicamycin (Tm). *P* values to respective WT or mutant worms on ND. *ns*, non-significant with *P*>0.05.

**Extended Data Fig. 5.**
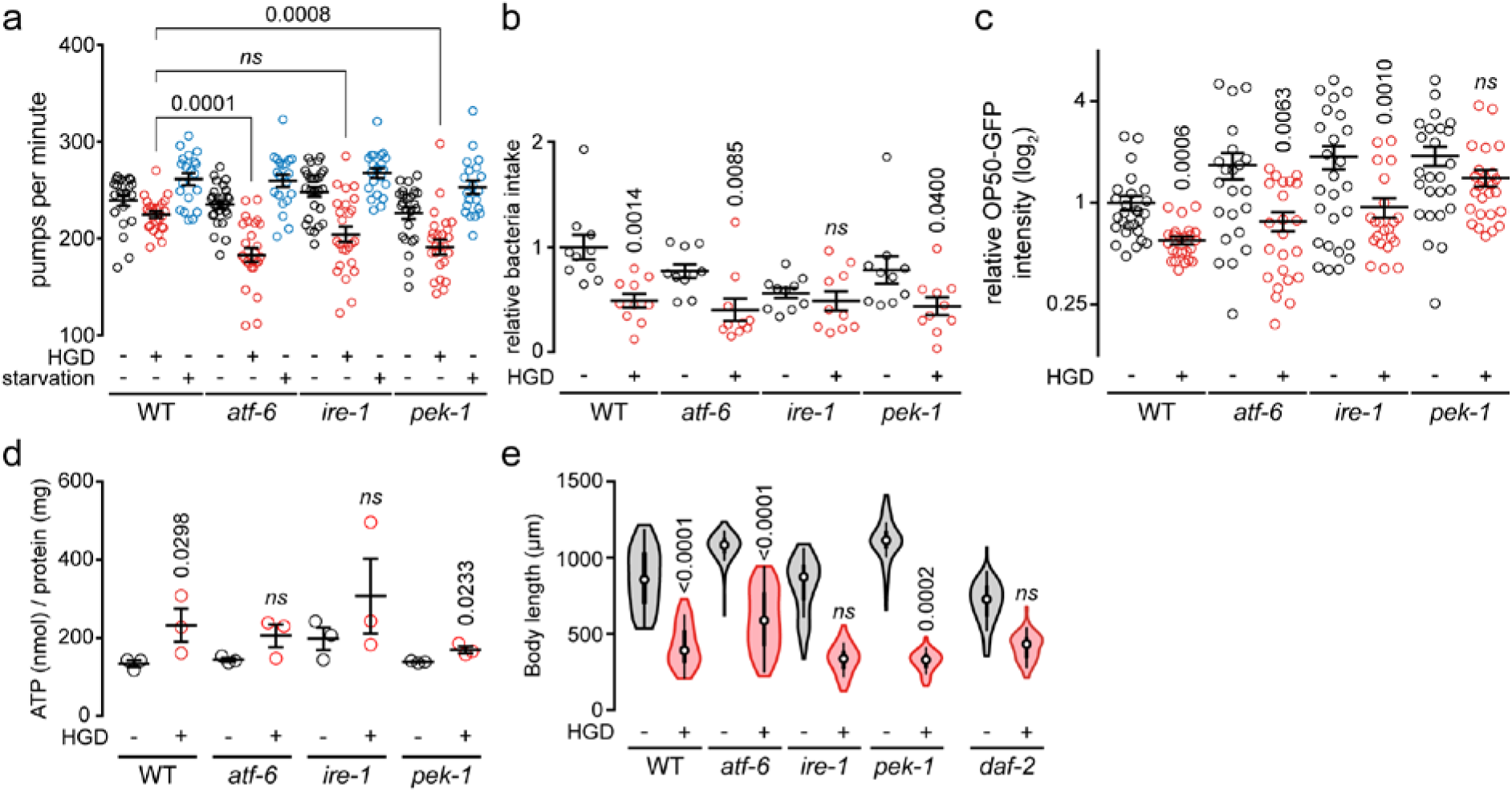
UPR mutant worms are metabolic active on high glucose diet. **a,** Pharyngeal pumping rate in 7-day-old WT and UPR mutants on 24h normal diet (ND), high glucose diet (HGD), or 4h starvation (n=22, n=25, n=20 for WT ND, HGD, starvation, respectively; n=27, n=27, n=20 for *atf-6(lof)* ND, HGD, starvation, respectively; n=27, n=25, n=21 for *ire-1(lof)* ND, HGD, starvation, respectively; n=25, n=26, n=20 for *pek-1(lof)* ND, HGD, starvation, respectively). **b,** Bacteria intake expressed relative to bacterial clearance within the WT and UPR mutants control interval of 7-day-old worms (n=46, n=39 for wild type worms on ND, HGD respectively; n=54, n=66 for *atf-6* mutant worms on ND, HGD respectively; n=67, n=62 for *ire-1* mutant on ND, HGD respectively; n=57, n=44 for *pek-1* mutant worm on ND, HGD respectively). **c,** Quantification of confocal fluorescent microscopy images of GFP-expressing bacteria colonising WT and UPR mutant worm intestine treated as in **a** (n=27, n=24 for WT ND, HGD, respectively; n=22, n=23 for *atf-6(lof)* ND, HGD, respectively; n=26, n=22 for *ire-1(lof)* ND, HGD, respectively; n=25, n=26 for *pek-1(lof)* ND, HGD, respectively). **d,** Total ATP concentration of worms treated as in **a**. **e,** Body length of worms treated as in **a**. *P* values to respective WT or mutant worms on normal diet. *ns*, non-significant with *P*>0.05.

**Extended Data Table 1.**
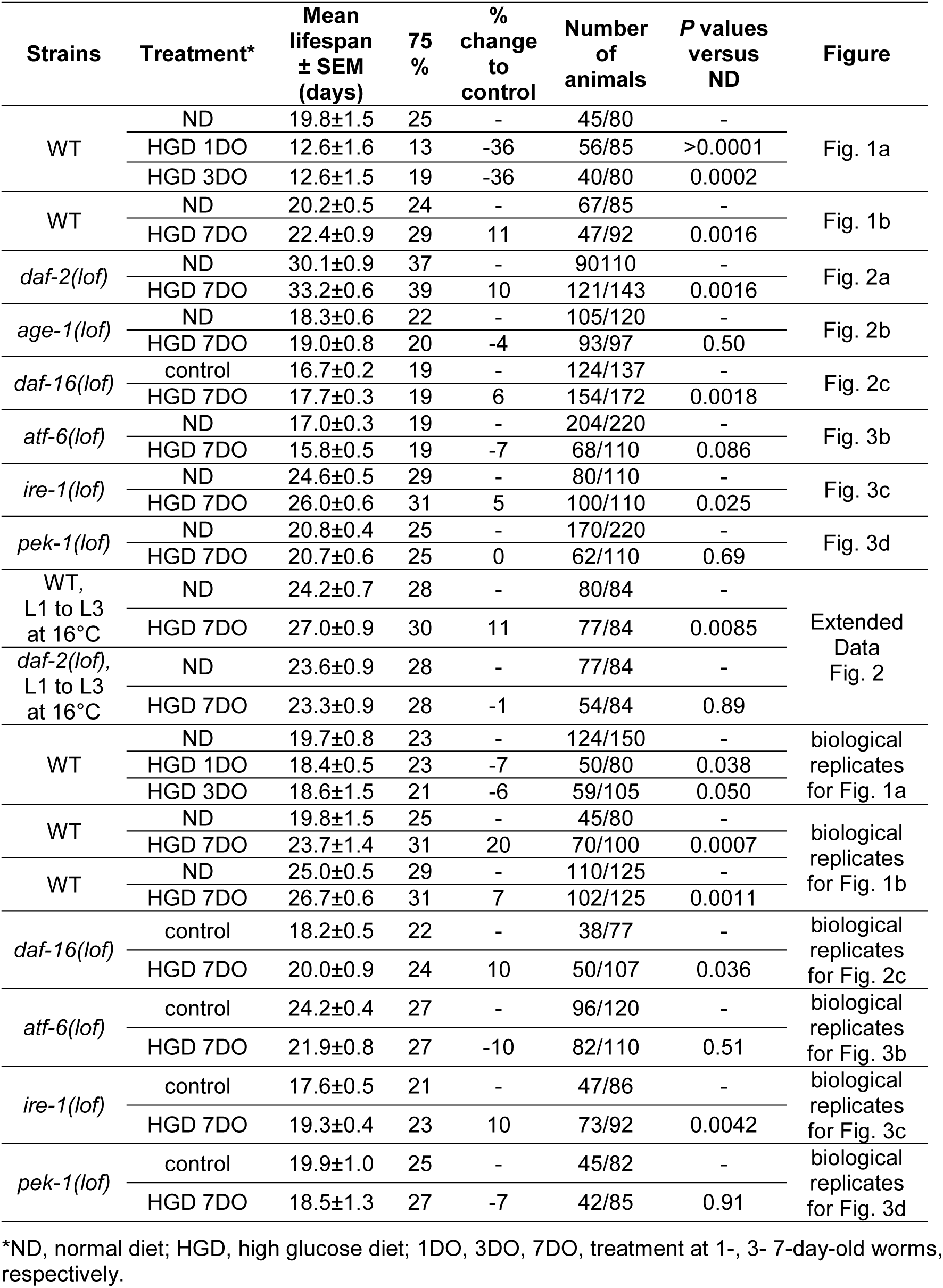
Lifespan analysis.

**Extended Data Table 2. UPR-upregulated genes modulating glucose inducible *far3p::GFP*. Excel Spreadsheet**

**Extended Data Table 3. Tissue-specific predominant GO terms. Excel Spreadsheet**

